# ID-Seg: An Accurate and Reliable Infant Deep learning Segmentation Framework for Limbic Structures

**DOI:** 10.1101/2021.03.29.437045

**Authors:** Yun Wang, Fateme Sadat Haghpanah, Xuzhe Zhang, Katie Santamaria, Gabriela Koch da Costa Aguiar Alves, Elizabeth Bruno, Natalie Aw, Alexis Maddocks, Cristiane S Duarte, Catherine Monk, Andrew Laine, Jonathan Posner, on behalf of program collaborators for Environmental influences on Child Health Outcomes

**Affiliations:** Department of Psychiatry, Columbia University Irving Medical Center, New York, NY, USA; New York State Psychiatric Institute, New York, NY, USA; Department of Biomedical Engineering, Columbia University, New York City, USA; Department of Computer Science, University Of Toronto, Toronto, Ontario, Canada; Department of Psychology, Columbia University, New York, NY, USA; Department of Obstetrics and Gynecology, Columbia University, New York, NY, USA; Department of Radiology, Columbia University, New York, NY,USA

## Abstract

Early postnatal period brain magnetic resonance imaging (MRI) is becoming an important approach to measure the impact of prenatal exposures on neurodevelopment and to investigate early biomarkers for risk. Among brain structures, Limbic structures are particular of interest in psychiatric disorder-related research. However, despite the promise of infant neuroimaging and the success of initial infant MRI studies, assessing limbic regions’ structure and function remains a significant challenge due to low inter-regional intensity contrast and high curvature (e.g., hippocampus). In addition, the agreement between existing automatic techniques and manual segmentation remains either untested or insufficient, particularly for the amygdala and hippocampus. In this work, we developed an accurate (based on three segmentation evaluation metrics), reliable and efficient infant deep learning segmentation framework (ID-Seg) to address the aforementioned challenges. Specifically, we leveraged a large dataset of 473 infant MRI scans to train ID-Seg and rigorously evaluated ID-Seg’s performance on internal and external datasets with manual segmentations. Compared with a state-of-the-art segmentation pipeline, we demonstrated that ID-Seg significantly improved the segmentation accuracy of limbic structures (hippocampus and amygdala) in newborn infants. Moreover, in a medium-size dataset, we found that ID-Seg-derived morphometric measures yield strong brain-behavior associations. As such, our ID-Seg may improve our capacity and efficiency to measure MRI-based brain features relevant to neuropsychological development and ultimately advance the success of quantitative analyses on large-scale datasets.

## 1. Introduction

Identifying early neurobiological markers of psychiatric risk is a critical step toward developing targeted early intervention strategies. Toward this goal, early postnatal period brain imaging studies are becoming increasingly common (Duarte, Monk, Weissman, & Posner, 2020; Posner et al., 2016). Adopting an early brain imaging approach 1) offers non-invasive and high-resolution images of infant brains, 2) opens up the potential to detect early biomarkers for risk, and 3) characterizes the infant brain with limited postnatal influence, and thus may help isolate, for instance, the impact of prenatal exposures on neurodevelopment. However, an important stumbling block curtailing progress in infant neuroimaging is developing accurate, reliable, and time-effective methods for segmenting brain regions on anatomical infant MRI scans.

Two candidate brain regions of particular interest for identifying psychiatric risk in infants are the amygdala and hippocampus. Studies of mood and anxiety disorders, but also of externalizing disorders, have repeatedly implicated altered amygdala and hippocampus structure and function. Infant studies have also suggested this relationship between structure and function (Graham et al., 2019; Lugo-Candelas et al., 2018; Qiu et al., 2013; Ramphal et al., 2020; Rogers et al., 2017). For example, amygdala-prefrontal functional connectivity in neonates is associated with internalizing symptoms at age 2 (Rogers et al., 2017); prenatal maternal depression is associated with altered white matter connectivity between the amygdala and prefrontal cortex on infant diffusion MRI scans (Lugo-Candelas et al., 2018); and maternal anxiety slows the development of hippocampus over the first 6 months of life (Qiu et al., 2013).

Despite the promise of infant neuroimaging and the success of initial infant MRI studies, assessing amygdala and hippocampus structure and function remains a significant challenge. First, both structures have relatively small volumes, meaning even small segmentation errors can lead to substantial miscalculations of morphometric estimates. Second, both structures (particularly the hippocampus) have a high degree of curvature, making it difficult for automated segmentation to accurately delineate these structures from surrounding neural tissue. Third, the amygdala and hippocampus are adjacent structures with low inter-regional tissue contrast, making it difficult to discern when one structure ends, and the other begins. As a result of these challenges, the accuracy of existing automated segmentation techniques remains limited. For example, although automatic segmentation pipelines have been developed (Dai, Shi, Wang, Wu, & Shen, 2013; Makropoulos et al., 2018; Zollei, Iglesias, Ou, Grant, & Fischl, 2020), the agreement between these automatic techniques and expert manual segmentation remains either untested or poor particularly for the amygdala and hippocampus, e.g., 0.39 agreement between automatic and manual segmentation *(Hashempour et al*., *2019)*. This scenario is further compounded when researchers apply automated techniques to datasets acquired from different MRI scanners or imaging protocols. Accurate, reliable, and time-efficient segmentation of the infant’s brain is important for advancing the success of virtually all quantitative analyses across MRI modalities.

This study aimed to develop and validate an accurate and reliable hippocampus and amygdala segmentation method using deep learning and transfer-learning methods for infant MRI research, henceforth termed “**ID-Seg**” (**I**nfant **D**eep learning **SEG**mentation). We began by initially training a 3-dimensional **ID-Seg** model on a publicly available neonatal structural MRI dataset (n=473) with brain regions segmented and labeled using an automated pipeline. Then, leveraging transfer-learning, we fine-tuned and internally tested our model on a smaller infant MRI dataset (n=20) with the amygdala and hippocampus manually segmented by three trained raters. To test the robustness of our **ID-Seg** model, we also included an external, publicly available dataset (n=10) with expert manual segmentations of the amygdala and hippocampus. We then compared the segmentation performance of **ID-Seg** to that of an existing automated pipeline, the Developmental human connectome [dHCP] pipeline (Makropoulos et al., 2018). Lastly, as an indicator of predictive validity, we examined prospective associations between a) amygdala and hippocampus morphometric measures from infant MRI scans (n=33) and b) parent-reported behavioral problems at age 2 as indexed by Child Behavior Checklist (CBCL) (Achenbach & Rescorla, 2000). We compared the magnitude of these associations based on morphometric measures derived from ID-Seg relative to those derived from dHCP. We hypothesized that **ID-Seg**-derived morphometric measures would provide stronger brain-behavior associations.

## 2. Materials and Methods

This study contains two modules. First, we developed and validated an infant deep learning segmentation framework (**ID-Seg**) for segmenting the infant hippocampus and amygdala using T2 weighted (T2w) MRI brain scans (**Figure 1.a**). Second, after training and optimizing our model, we directly applied optimized ID-Seg to segment a new input of infant MRI scans and explored prospective associations between brain structure at birth and behavioral problems at age 2 (**Figure 1.b**).

**Figure 1.**
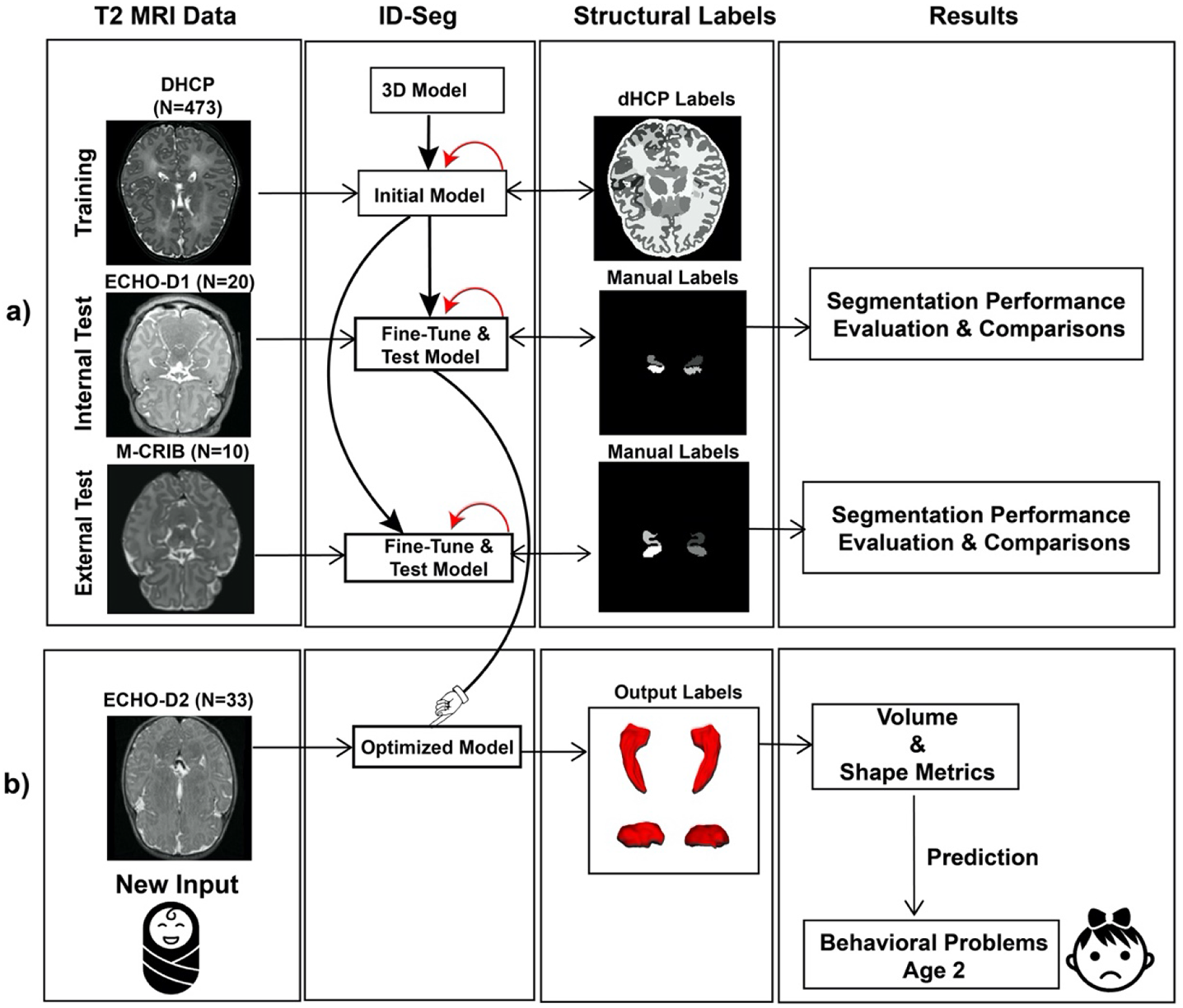
Overview of this study. **a)** We trained, fine-tuned, tested internally, and tested externally our deep-learning (DL) segmentation framework (ID-Seg) for newborn hippocampus and amygdala on three independent datasets; **b)** Using the optimized model, we directly segmented another batch of infants’ brain scans without expert segmentation and further explored the prospective associations **between** morphometric measures (L/R hippocampus and Amygdala) at birth and behavior problems at age 2. The red circle indicates parameters of ID-Seg were updated.

### 2.1 Datasets

To test the segmentation performance and reliability of ID-Seg on datasets from different scanners and manual raters, we curated 4 infant MRI datasets for initial model training, internal testing, external testing, and exploration of prospective associations. We provided demographics and imaging parameters for each of these four datasets in **Table 1**.

**Table 1.** Demographics and MRI Sequence Information.

| Table 1. Demographics and MRI Sequence Information |  |  |  |  |
| --- | --- | --- | --- | --- |
|  | Training DHCP*<br>(N=473) | ECHO-D1<sup>f</sup><br>(N=20) | M-CRIB<sup>†</sup><br>(N=10) | ECHO-D2<sup>‡</sup><br>(N=33) |
| PMA at scan, weeks | 40.65 ± 2.19 | 46.90 ± 4.14 | 39.78 ± 1.31 | 47.37 ± 5.01 |
| Sex |  |  |  |  |
| Female, N(%) | 266 (43.8%) | 13 (65.0%) | 4 (40.0%) | 14 (42.4%) |
| Male, N (%) | 207 (56.2%) | 7 (35.0%) | 6 (60.0%) | 19 (57.6%) |
| MRI scanners | 3T Philips | 3T GE | 3T Siemens | 3T GE |
| MRI resolution (mm<sup>3</sup>) | [0.5, 0.5, 0.5] | [0.9, 0.9, 0.9] | [0.63, 0.63, 0.63] | [0.9, 0.9, 0.9] |
| MRI Dimensions | [290,290,203] | [130,256,256] | [304,304,157] | [130, 256, 256] |

Note. — For quantitative variables, data are presented as mean ± standard deviation unless otherwise noted. PMA: postmenstrual age. \*The large **Training DHCP** dataset with corresponding dHCP labels was used to pre-train the model with sufficient data. <sup>f</sup>The **ECHO-D1** dataset was used to test the model's performance. <sup>†</sup>The **M-CRIB** dataset was used as an external test dataset to further test the reliability of our proposed deep learning framework and output an optimized ID-Seg model. <sup>‡</sup>The **ECHO-D2** dataset contains corresponding CBCL measures at age 2-year-old. It is used to compare the predictive validity between segmentation results from dHCP pipeline and ID-Seg.

#### Training Dataset – Developmental human connectome (dHCP) project

For our training dataset, we included **473** bias-corrected infant T2-weighted (T2w) structural MRI scans from the dHCP v1.0.2 data release (note: we excluded 33 T2w scans acquired in a sagittal plane, different from the 473 scans acquired in an axial plane). Bilateral hippocampus and amygdala dHCP labels were generated by the dHCP structural segmentation pipeline (Makropoulos et al., 2018). Interested readers can find additional details about the imaging parameters and preprocessing procedures in the original work (Makropoulos et al., 2018).

#### Internal testing dataset – Environmental influences on child health outcomes (ECHO)-D1

For our internal test dataset, we used **20** term-born high-quality infant T2w MRI scans. These 20 scans were randomly drawn from the following two ECHO studies – 10 were chosen from ECHO– Boricua Youth Study at the New York State Psychiatric Institute (ECHO-BYS-NYSPI) and 10 from ECHO-BYS at the University of Puerto Rico (ECHO-BYS-UPR). Detailed descriptions about the two studies, including subject enrollment, imaging parameters, and inclusion/exclusion criteria, can be found in ***Supplemental Materials: Dataset***. In addition, we manually segmented the bilateral hippocampal and amygdaloid structures using a multi-rater method (see section 2.2).

#### External testing dataset – Melbourne Children’s Regional Infant Brain (M-CRIB)

For our external test dataset, we used a publicly available dataset including T2w MRI scans of 10 term-born infants with manual segmentations performed by a single rater from an independent group (Alexander et al., 2017). Detailed descriptions about the manual segmentation procedures and imaging parameters can be found elsewhere (Alexander et al., 2017).

#### Prospective associations – ECHO-D2

As a proof-of-concept analysis, we explored prospective associations in an additional 33 infants. These infants had both T2w MRI scans and CBCL assessments completed as part of the aforementioned ECHO-BYS-NSYPI and ECHO-BYS-UPR studies (these infant data do not overlap with those of ECHO-D1). We obtained these MRI scans at ∼1-4 months of age and subsequent parent-report CBCL assessments at ∼2 years of age.

### 2.2 Multi-rater manual segmentatio

Three research assistants received instruction and training from a board-certified radiologist to perform infant hippocampus and amygdala segmentation using ITK-SNAP software (Yushkevich et al., 2006). Manual segmentation protocols and inter-rater reliability assessments are available in the **Supplemental Materials**. For each brain region, the manual tracing’s inter-rater reliability was assessed and ensured before proceeding (a minimum 0.6 dice similarity coefficient). Based on all three raters’ manual segmentation, a bilateral reference manual segmentation for the amygdala and hippocampus was generated with the Simultaneous Truth And Performance Level Estimation (STAPLE) algorithm (Warfield, Zou, & Wells, 2004). STAPLE is an expectation-maximization algorithm that estimates the optimal combination of segmentation based on each rater’s performance level. We assessed the “ground truth” segmentation by the STAPLE based on all three study raters instead of only one rater.

### 2.3 Infant Deep Learning Segmentation (ID-Seg)

We adopted a transfer-learning approach to train and test ID-Seg on multiple datasets. In **Figure 1.a**, ID-Seg was initially trained (termed “pre-train” in the AI literature) on our training dataset, **dHCP**, consisting of 473 participants’ T2w MRI scans. Then we tested this trained model on internal and external datasets – **ECHO-D1** and **M-CRIB**, respectively. For the ECHO-D1 dataset, we used our multi-rater manual segmentation framework as described above to generate manual segmentations; for M-CRIB, researchers from an independent group (Alexander et al., 2017) generated manual segmentations and made these publicly available (https://osf.io/4vthr/). Finally, to evaluate model performance, we calculated the dice similarity coefficients (DSC) and average surface distance (ASD) between the manual segmentation and the predicted labels from our deep learning model (see section 2.4). All deep learning models below were written in Python using PyTorch libraries, and relative training procedures were completed in the NVIDIA Geforce Titan RTX GPU workstation. In addition, we shared relevant codes in an open-access GitHub repository (https://github.com/wangyuncolumbia/ID-Seg).

#### 2.3.1 Model Architecture

We used a multi-view fully convolutional neural network (F-CNN) for 3D hippocampus and amygdala segmentation. This model was initially developed for adult whole brain segmentation (Guha Roy, Conjeti, Navab, Wachinger, & Alzheimer’s Disease Neuroimaging, 2019) and has been demonstrated to be capable of segmenting small subcortical structures with skip connections and unpooling layers in the decoding path. Specifically, we trained three 2D F-CNN models separately for each of the three principal views (axial, coronal, and sagittal). Of note, each 2D F-CNN has the same architecture. In the end, we merged predicted probabilities from multi-view models using formula (1) below to calculate the final predicted label for each voxel.

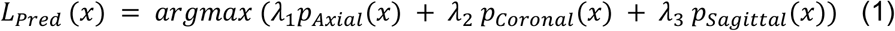

In formula (1), *p*_*Axial*_ (*x*), *p*_*Coronal*_ (*x*), *p*_*Sagittal*_ (*x*) are predicted probabilities of a voxel from axial, coronal, and sagittal deep learning models. We set the weights *λ*_1_, *λ*_2_, *λ*_3_ to 0.4, 0.4, and 0.2, respectively. Additional details about this model can be found here (Guha Roy et al., 2019).

#### 2.3.2 Model Learning

##### 2.3.2a Initial network training

The goal of training ID-Seg on a large **dHCP** dataset first was to provide robust weight initialization. We randomly split this training dataset into two parts: 80% for training and 20% for validating model performance. As noted above, we applied the dHCP structural segmentation pipeline, which bilaterally segments and labels 87 regions within the infant’s brain, including the hippocampus and amygdala. We started with these automated segmentations because we reasoned that this large sample would provide strong prior initialization of the network, such that we could then optimally utilized the smaller sample of manually segmented scans to achieve high segmentation accuracy. We anticipated that the segmentations from the automated software (e.g., dHCP) would not be as accurate as the manual annotations. However, these segmentations would allow our model to recognize a wide range of morphological variations in brain structures. This training procedure affords strong prior weights for the ID-Seg network, where robustness to data heterogeneity is enhanced by the diversity of the training dataset (e.g., different scanners and sites).

##### 2.3.2b Internal testing

We next applied the initially trained ID-Seg and fine-tuned it on **ECHO-D1** (see **Section 2.2, Figure 1.a**). Specifically, we first passed weights of the initially trained ID-Seg, and we then trained ID-Seg for 5 epochs while only unfreezing the last few layers to prevent propagation errors due to random initialization weights and save computation time. Lastly, we unfroze all layers and fine-tuned the whole network for another 50 epochs. Similarly, we evaluated ID-Seg’s performance against that of the manual segmentation we conducted on the internal datasets by a multi-rater framework using the leave-one-out cross-validation (LOOCV) technique. The hyperparameters used to fine-tune the network, including learning rate, batch size, and epoch number, are 5*10^−4^, 8, and 50, respectively.

##### 2.3.2c External testing

To test the reliability of ID-Seg, similarly, we applied ID-Seg to a publicly available dataset, **M-CRIB** (see section 2.1), and evaluated the accuracy of ID-Seg’s segmentations against manual segmentations performed by an independent group. Again, we used similar hyperparameters as in internal testing.

### 2.4 Evaluation metrics

We calculated three commonly used evaluation metrics to compare the segmentation output of our ID-Seg against manual segmentations: Dice Similarity Coefficient (DSC), intra-class correlation (ICC), and Average Surface Distance (ASD). DSC is a metric used to calculate the similarity between two images and measure the overlap across the two images (Dice, 1945). ICC is a measure of consistency between two raters, or in this case, two segmented images (Shrout & Fleiss, 1979). ASD is a surface-based metric and measures the average Hausdorff Distance over all points between surfaces of a prediction structure (i.e., ID-Seg’s segmentation of the hippocampus and amygdala) and the “‘ground truth”’ (i.e., manually segmented hippocampus and amygdala). (the code can be found at https://github.com/deepmind/surface-distance). Of note, there are ∼20 different metrics to evaluate segmentation. Readers can find a throughout review elsewhere (Taha & Hanbury, 2015).

### 2.5 Volume and Shape analysis

We applied the optimized version ID-Seg to directly segment infant scans in the **ECHO-D2** dataset – that is, infant MRI scans that were not used in any of the previous training/testing procedures (**Figure 1.a**). Using ID-Seg, we calculated volumetric and shape measurements for each of four structures (bilateral hippocampus and amygdala). The volume (in mm^3^) of each region was calculated based on pixel numbers and pixel dimension in the segmented binary mask. Then, we performed shape analysis for each structure using SlicerSALT software (Kitware, Inc., United States). Here, we used an average spherical harmonics description (SPHARM) to represent the shape measurements of a 3D structure (Styner et al., 2006). For each structure (L/R hippocampus and amygdala), we derived one volume and one shape measurement.

### 2.6 Statistical analysis

To compare segmentation metrics derived from **ID-Seg** relative to those from the dHCP pipeline, we conducted paired t-tests for volume and shape measures derived above (see section 2.5). We also performed FDR correction to adjust estimated p values for multiple comparisons.

Further, we conducted Spearman rank partial correlation analysis to investigate prospective associations between volume and shape measures on infant MRI and behavioral outcomes at age 2. Behavior outcomes including internalizing, externalizing, and total problems were assessed using the parent-report CBCL. Postmenstrual age at MRI scan was controlled during the analysis. Sex was not adjusted for because this has already been accounted for in the CBCL T scores.

Lastly, we also adopted a multivariate approach – partial least square regression (PLSR) to predict CBCL scores at age 2. We used PLSR due to our relatively small sample size and multicollinearity among the 8 brain measures described in section 2.5 (see ***Supplemental Figure 1*** for multicollinearity test). We evaluated the generalization error of PLSR by minimizing root-mean-square error (RMSE) with the LOOCV technique. PLSR can detect latent features that simultaneously summarize the predictors’ variation while optimally correlating these predictors with the outcome of interest. We calculated cross-validated RMSE (RMSECV) and R^2^ to estimate the accuracy and generalization of the PLSR model. All statistical analyses were conducted in R.

## 3. Results

### 3.1 Segmentation Performance of ID-Seg relative to “Groud-Truth” Segmentations “Groud-Truth” Segmentations

For the internal test dataset ECHO-D1 (n=20), our multi-rater manual segmentation framework (see section 2.2) generated ground-truth segmentation of the bilateral hippocampus and amygdala; the overall inter-rater agreement across all structures was **0.76 (0.06)**. For each structure, the mean and standard deviation were 0.78 (0.05) for R hippocampus, 0.77 (0.05) for L hippocampus, 0.73 (0.05) for R amygdala, and 0.74 (0.06) for L amydala. Additional quantitative and segmentation protocols are in our ***Supplemental Table 1*** and ***Supplemental Materials*** “*Manual Segmentation Protocols using ITK-SNAP: Hippocampus and Amygdala*.”

For each dataset, we next evaluated the accuracy of ID-Seg-derived and dHCP-derived segmentations against these “ground-truth” labels using two metrics, DSC and ASD (see section 2.4).

#### Segmentation Metrics of ID-Seg

For the ECHO-D1 dataset, the average DSC of ID-Seg across four structures was 0.86 (0.03); ICC was 0.93 (0.02);ASD was 0.29 (0.11) mm. For the the M-CRIB dataset, ID-Seg’s performance was similar: 0.87 (0.02) for DSC, 0.93 (0.01) for ICC, and 0.32 (0.10) for ASD. Each measure’s mean and standard deviation were reported in **Table 2**, and the distribution is in **Figure 2**.

**Figure 2.**
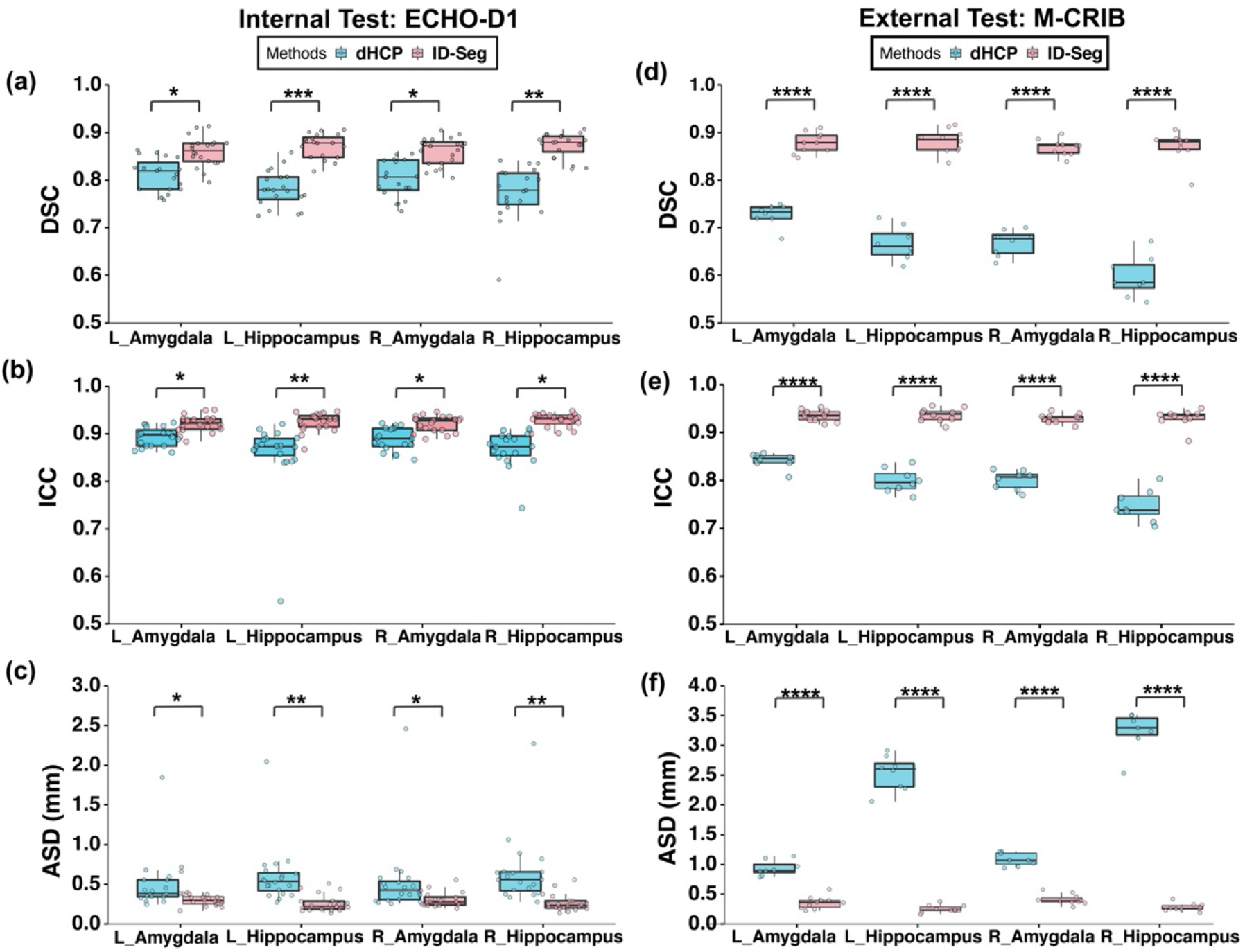
ID-Seg significantly improved the segmentation accuracies on two independent infant datasets and are comparable with expert manual segmentations. Panels **a), b) and c)** illustrated segmentation performance on the ECHO-D1 dataset using the Dice Similarity Coefficients (DSC), intra-class correlation (ICC) and average surface distance (ASD), respectively; Panel **d), e)** and **f)** illustrated segmentation performance on M-CRIB dataset using DSC, ICC and ASD. We also compared the segmentation of ID-Seg with that of dHCP pipeline. * denotes p<0.05, ** denotes p<0.01, *** denotes p<0.001, **** p<0.0001

**Table 2.** Segmentation Metrics and Comparisons between dHCP and ID-Seg.

| Table 2. Segmentation Metrics and Comparisons between dHCP and ID-Seg |  |  |  |  |  |  |  |
| --- | --- | --- | --- | --- | --- | --- | --- |
| Datasets | Metrics | Regions | Methods |  | statistic | p | p.adj |
|  |  |  | dHCP | ID-Seg |  |  |  |
| **ECHO-D1**<br>(n=20) | **DSC** | L Amygdala | 0.79 (0.12) | 0.86 (0.03) | -2.69 | 0.014 | 0.014 |
|  |  | L Hippocampus | 0.76 (0.09) | 0.87 (0.03) | -4.506 | <10<sup>-3</sup> | 0.001 |
|  |  | R Amygdala | 0.77 (0.14) | 0.86 (0.03) | -2.69 | 0.014 | 0.014 |
|  |  | R Hippocampus | 0.74 (0.15) | 0.87 (0.03) | -3.911 | 0.001 | 0.002 |
|  | **ICC** | L Amygdala | 0.87(0.09) | 0.92 (0.02) | -2.252 | 0.035 | 0.044 |
|  |  | L Hippocampus | 0.86 (0.08) | 0.93 (0.02) | -3.751 | 0.001 | 0.005 |
|  |  | R Amygdala | 0.86 (0.13) | 0.92 (0.02) | -2.153 | 0.044 | 0.044 |
|  |  | R Hippocampus | 0.84 (0.13) | 0.93 (0.01) | -2.974 | 0.008 | 0.015 |
|  | **ASD** | L Amygdala | 0.49 (0.34) | 0.32 (0.11) | 2.202 | 0.038 | 0.05 |
|  |  | L Hippocampus | 0.60 (0.37) | 0.26 (0.11) | 3.939 | 0.001 | 0.002 |
|  |  | R Amygdala | 0.53 (0.47) | 0.31 (0.09) | 2.085 | 0.05 | 0.05 |
|  |  | R Hippocampus | 0.65 (0.43) | 0.26 (0.11) | 3.916 | 0.001 | 0.002 |
| **M-CRIB**<br>(n=10) | **DSC** | L Amygdala | 0.73 (0.02) | 0.88 (0.02) | -14.124 | <10<sup>-4</sup> | <10<sup>-4</sup> |
|  |  | L Hippocampus | 0.67 (0.04) | 0.88 (0.03) | -14.73 | <10<sup>-4</sup> | <10<sup>-4</sup> |
|  |  | R Amygdala | 0.67 (0.03) | 0.87 (0.02) | -18.297 | <10<sup>-4</sup> | <10<sup>-4</sup> |
|  |  | R Hippocampus | 0.60 (0.04) | 0.87 (0.03) | -15.259 | <10<sup>-4</sup> | <10<sup>-4</sup> |
|  | **ICC** | L Amygdala | 0.86(0.05) | 0.93(0.01) | -13.673 | <10<sup>-4</sup> | <10<sup>-4</sup> |
|  |  | L Hippocampus | 0.83(0.06) | 0.94(0.01) | -13.927 | <10<sup>-4</sup> | <10<sup>-4</sup> |
|  |  | R Amygdala | 0.83(0.06) | 0.93(0.01) | -17.022 | <10<sup>-4</sup> | <10<sup>-4</sup> |
|  |  | R Hippocampus | 0.78(0.09) | 0.93(0.02) | -14.119 | <10<sup>-4</sup> | <10<sup>-4</sup> |
|  | **ASD** | L Amygdala | 0.94 (0.13) | 0.36 (0.11) | 10.268 | <10<sup>-4</sup> | <10<sup>-4</sup> |
|  |  | L Hippocampus | 2.5 (0.29) | 0.25 (0.06) | 21.747 | <10<sup>-4</sup> | <10<sup>-4</sup> |
|  |  | R Amygdala | 1.1 (0.11) | 0.41 (0.09) | 14.192 | <10<sup>-4</sup> | <10<sup>-4</sup> |
|  |  | R Hippocampus | 3.3 (0.44) | 0.28 (0.06) | 19.619 | <10<sup>-4</sup> | <10<sup>-4</sup> |

**DSC**, Dice Similarity Coefficients; **ICC**, intra-class correlation; **ASD**, Average Surface Distance, measured in mm. segmentation metrics for each method were shown in mean (standard deviation). Higher DSC and ICC, lower ASD indicate better segmentation performance.

#### Segmentation Metrics of dHCP pipeline

For the ECHO-D1 dataset, dHCP’s average DSC was 0.77 (0.13), average ICC was 0.86 (0.11) and dHCP’s average ASD was 0.57 (0.40). For the M-CRIB dataset, we noticed the performance of dHCP significantly dropped: 0.66 (0.06) for DSC, 0.79 (0.04) for ICC, and 2.0 (1.1) mm for ASD. Each feature’s mean and standard deviation can be found in **Table 2**.

#### ID-Seg vs. dHCP

On both internal and external datasets (ECHO-D1 and M-CRIB, respectively), ID-Seg performed significantly better than dHCP pipeline. Complete statistical results are available in **Table 2**. As shown in Figure 3, compared with the dHCP pipeline, we found that structures segmented by ID-Seg had: 1) smaller volumes; 2) smoother surfaces; and 3) the 3D shape is closer to morphometric features.

**Figure 3.**
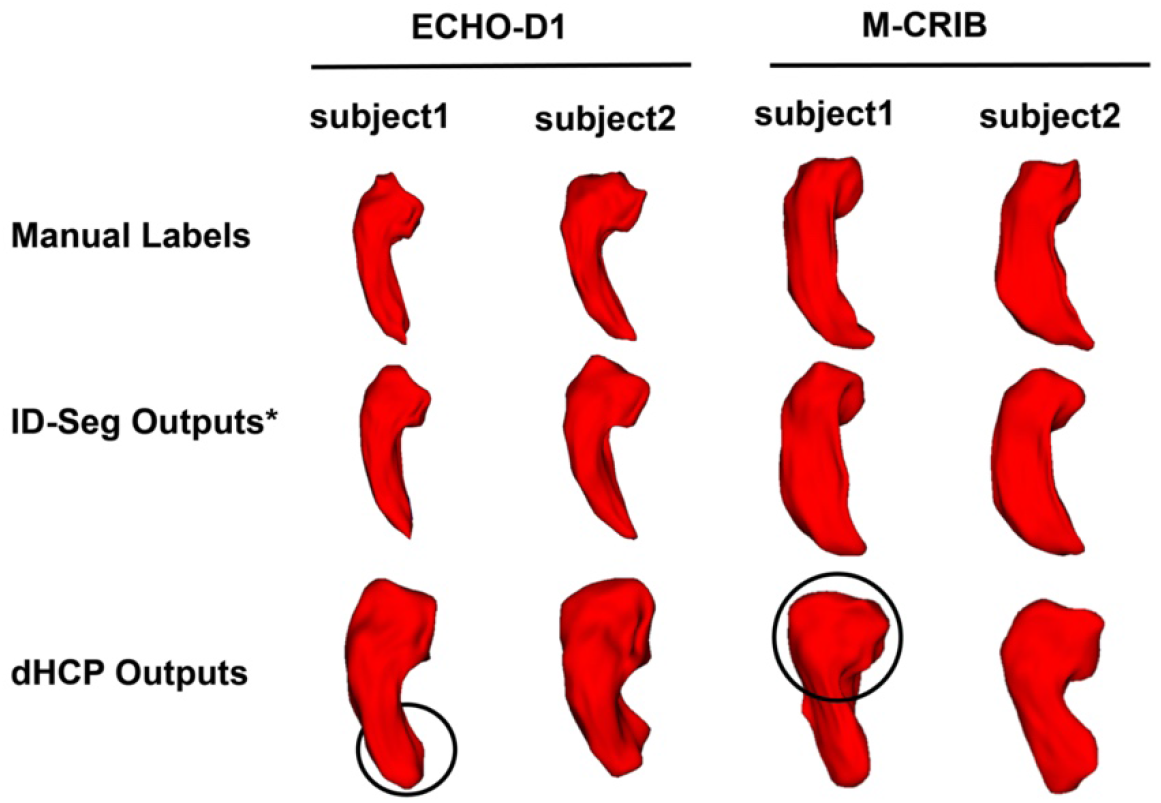
Visual comparisons among the “ground-truth” manual segmentation, ID-Seg segemntation outputs, and dHCP segementation outputs for left hippocampus using ECHO-D1 and M-CRIB dataset respectively. Compared with the dHCP pipeline, we found that structures segmented by ID-Seg had: 1) smaller volumes; 2) smoother surfaces ; and 3) the 3D output is closer to morphometric features (see hippocampus tail and head, black circle highlighted).

In sum, our results demonstrated good-to-excellent accuracy of ID-Seg when evaluated against “ground-truth” manual segmentations from different sources and scanners. Further, compared against the dHCP pipeline, ID-Seg improved the accuracy of hippocampus and amygdala segmentation.

### 3.2 Bivariate Analysis - Relationship between Brain at birth and CBCL Outcomes at 2

After directly applying ID-Seg to 33 infant scans from the ECHO-D2 dataset, we derived volume and shape metrics from the bilateral hippocampus and amygdala (see section 2.5). In addition to these brain features, we also obtained parent-reported behavioral problems when these infants matured to age 2, assessed by CBCL T scores in the following domains: internalizing, externalizing, and total problems. The mean and standard deviation of CBCL T scores for each domain were 49.1 (11.3) for total problems, 48.9 (11.4) for internalizing problems, 48.6 (9.7) for externalizing problems. We next investigated non-parametric relationships between brain and behavior.

As seen from **Figure 4.a**, in the analysis, we identified multiple significant correlations (13 out of 24) between ID-Seg derived brain features and age 2 behavioral outcomes. For example, we found significant correlations between the volume of right amygdala and CBCL total problems, rho(30) = -0.62, p<10^−3^; internalizing problems, rho(30) = -0.43, p<10^−3^; and externalizing problems, rho(30) = -0.59, p<10^−3^.

**Figure 4.**
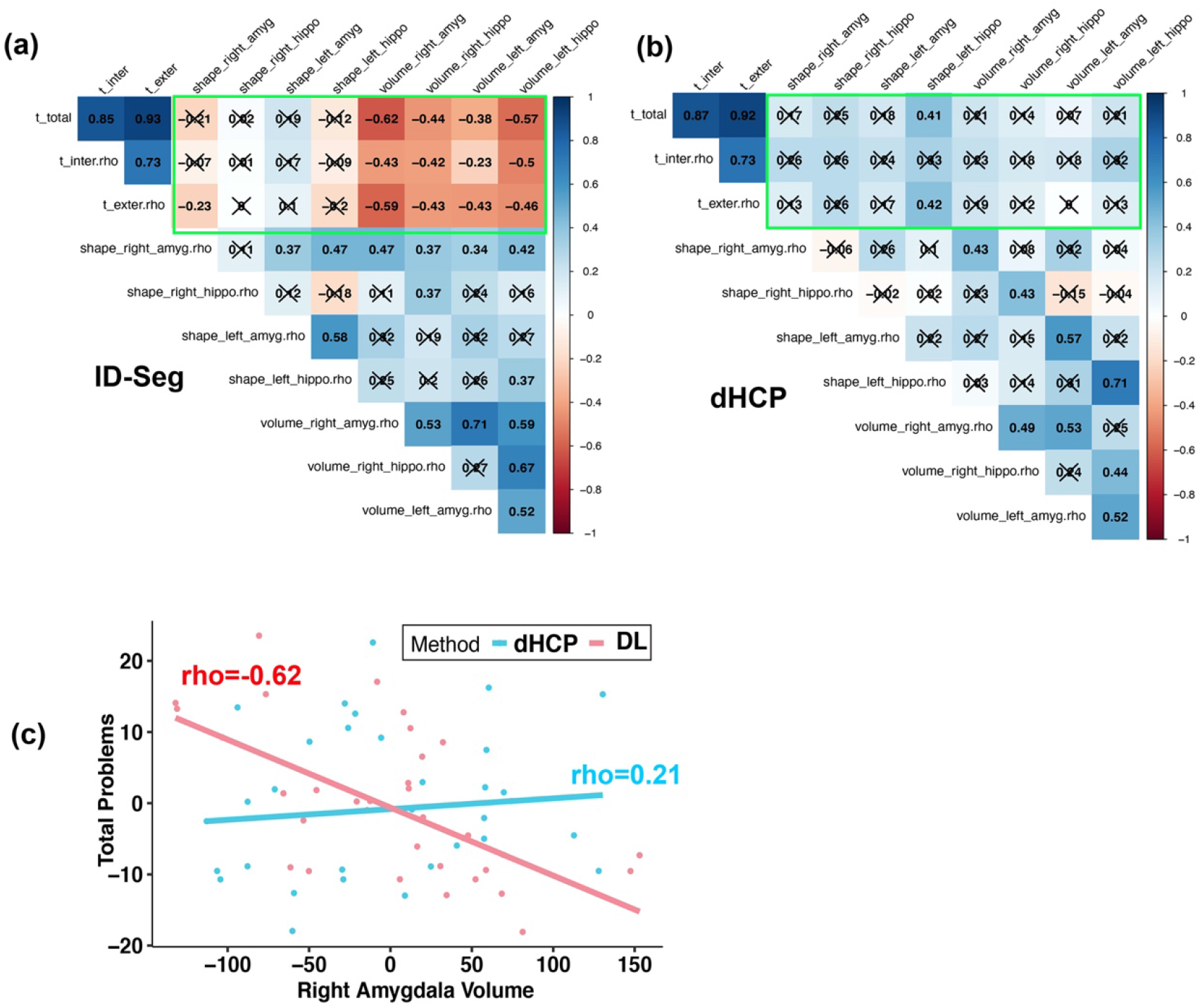
Pairwise non-parametric brain-behavior relationships. (in green rectangles) for two methods: **a**) our ID-Seg, **b)** dHCP. **X** indicates that the p value of spearman rank correlation is greater than 0.05. As an example, in **c)**, we created the scatterplot between age-adjusted CBCL total problems and age-adjusted right amygdala shape measures; the rho using dHCP method is 0.21, and in contrast, -0.62 using DL method.

Using dHCP-derived brain features, brain-behavior relationships are presented in **Figure 4.b** for comparison. Surprisingly, we only observed 2 out of 24 significant brain-behavior relationships. For example, the relationship between the volume of the right amygdala and each domain of the CBCL behavioral outcomes was not significant: total problems, rho(30)=0.21, p=0.46; internalizing problems, rho(30)=0.23, p=0.39; and externalizing problems, rho(30)=0.19, p=0.53. To visualize the contrast, we show pairs of age-adjusted total problems and age-adjusted right amygdaloid average volume and fitted regression lines in **Figure 4.c**.

### 3.3 Multivariate Analysis – Predicting Behavioral Problems at age 2

We generated three partial least square regression analysis models to predict each domain of behavioral problems at age 2: PLSR_total_, PLSR_internalizing_, and PLSR_externalizing_.

Using ID-Seg-derived brain features (**Figure 5.a**), two of three PLSR models were moderately predictive of behavioral problems at age 2. Specifically, in PLSR_total_ model (**Figure 5.b red line**), RMSECV and R^2^ values are 7.7 and 0.52, respectively; in PLSR_internalizing_ model (**Figure 5.c red line**), RMSECV and R^2^ values are 8.6 and 0.22, respectively; and in PLSR_externalizing_ model (**Figure 5.d red line**), RMSECV and R^2^ values are 8.8 and 0.40, respectively.

**Figure 5.**
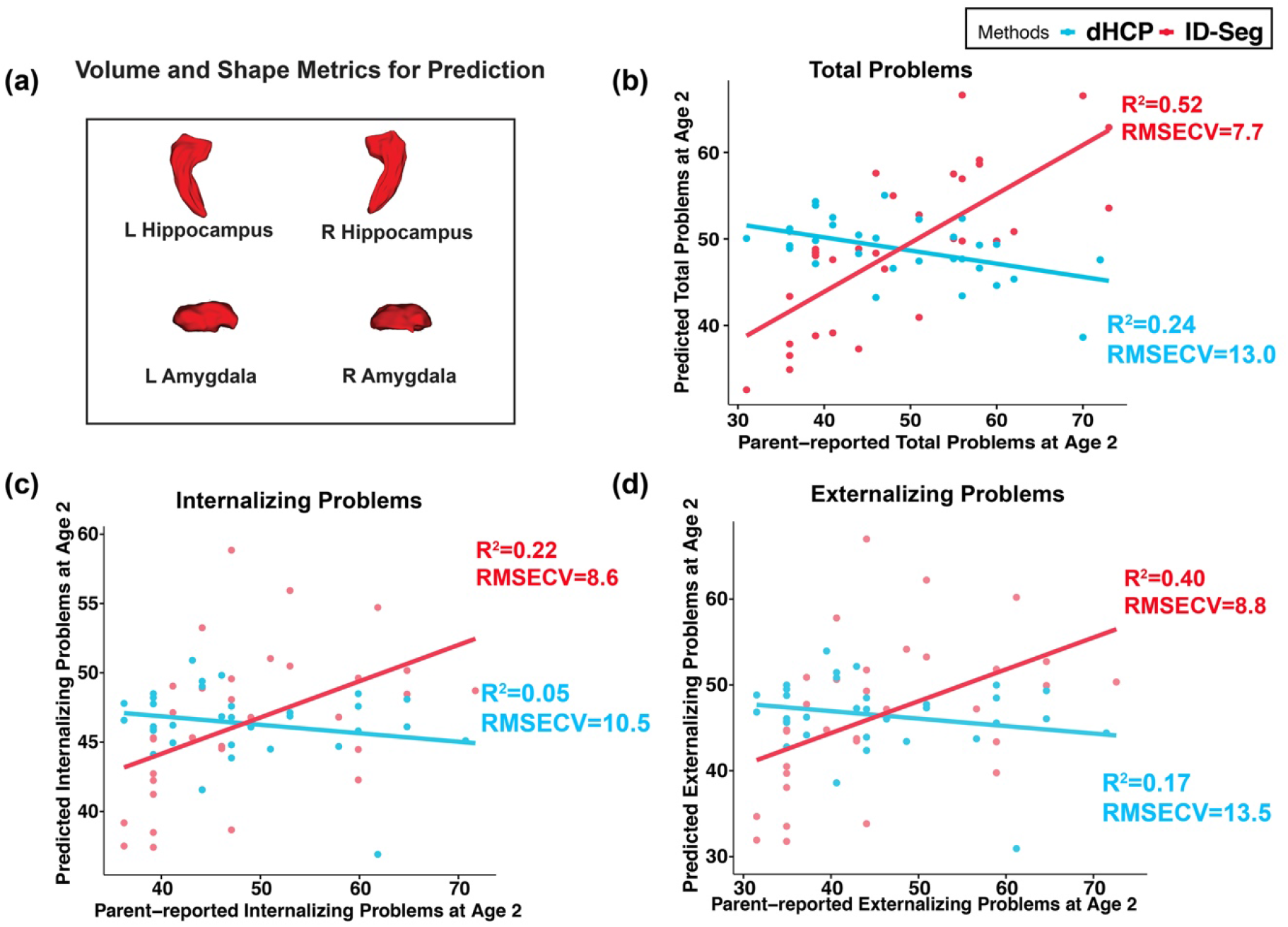
Using a mutlivariate approach (partial least square regression), hippocampal and amygdaloid measures derived by ID-Seg were collectively predictive of behavioral problems at age 2, as indicated by the **red lines** in **a)** CBCL total problems, **b)** CBCL internalizing problems, and **c)** CBCL externalizing problems. In contrast, dHCP-derived measures performed poorly for prediction, as indicated by the **blue lines** in panels **a), b)** and **c)**.

In contrast, using dHCP-derived brain features, all three PLSR models only showed marginal associations with CBCL total, internalizing, and externalizing problems (see **Figure 5.b, 4.c, 4.d blue lines)**.

Furthermore, we used the variable importance in the projection (VIP) method to compute each predictor’s influence on the outcomes. Consistently, we found that the right amygdala’s volume is the most salient brain feature in predicting behavioral outcomes.

## 4 Discussion

Our study used a large dataset of 473 infant MRI scans to train a 3D deep learning infant segmentation model (ID-Seg) and then evaluated ID-Seg’s performance on internal (ECHO-D1, n=20) and external datasets (M-CRIB, n=10) with manual segmentations of the hippocampus and amygdala. We found that: 1) ID-Seg achieved a high degree of segmentation accuracy: an average of 0.86 and 0.87 DSC, an average of 0.93 and 0.93 ICC, and an average of 0.29 and 0.32 ASD on the internal and external datasets, respectively; and 2) ID-Seg’s hippocampal and amygdaloidal estimates were more accurate than those obtained from dHCP (a commonly used pipeline).

We also explored prospective associations in another dataset (ECHO-D2, n=33), testing whether the volumes and shapes of the hippocampus and amygdala estimated from ID-Seg were associated with parent-reported behavioral problems in these same toddlers at age 2. We found that the volumes and shapes estimated by ID-Seg correlated with behavioral problems with moderate effect sizes. Conversely, using the same dataset, volumetric and shape estimates of the same structures from the dHCP pipeline demonstrated non-significant correlations with behavioral problems. Additionally, using a multivariate approach (PLSR), ID-Seg derived hippocampal and amygdaloidal measures were collectively predictive of behavioral problems at age 2 (R^2^ = 0.52, 0.22, and 0.40, respectively). In contrast, dHCP-derived measures performed poorly (R^2^ = 0.24, 0.05, and 0.17, respectively).

An accurate, computationally efficient, and reliable method for segmenting structural MRI scans is an important step in human neuroimaging research. This need is particularly evident in the current research world where large-scale, multi-site studies are increasingly common – studies such as the Adolescent Brain Cognitive Development (ABCD) study that includes over 10,000 MRI scans and involves multiple research sites with inter-site differences in MRI scanners. Whereas progress toward this goal has been made for studies of adults and youth, structural MRI scans of infants require special consideration because of the marked differences in tissue contrast seen in the infant relative to adult or child brain. For example, imaging pipelines such as FreeSurfer that are widely used in adult and youth studies, including the ABCD study, are not able to segment common subcortical structures on infant structural MRI scans. To our knowledge, only two MRI processing pipelines have been developed for infant subcortical segmentation: infant FreeSurfer and dHCP. Both offer important advances to the field, but both also have important limitations. The Infant FreeSurfer pipeline, in keeping with its adult counterpart, is optimized for T1w scans, yet most structural MRI studies in infant research use T2w scans because they provide clearer grey-white boundaries in the infant’s brain. Thus, we did not compare ID-Seg with infant Freesurfer because infant Freesurfer requires T1w scans as input. We suspect two potential reasons for why dHCP did not perform as well as ID-Seg in segmenting limbic structures. First, the dHCP segmentation pipeline was developed based on 20 manually segmented infant brain scans; however, 15 of these were from infants born prematurely. Samples used in our work are born from full-term infants, and numerous previous studies have shown significant structural brain differences in pre-relative to full-term infants (Ment & Vohr, 2008; Ophelders et al., 2020). Second, MRI scans in our work were collected from multiple vendors, including GE, Siemens, and Philips, allowing ID-Seg to learn and adapt to the idiosyncratic features of specific MRI vendors. Scanner differences might have caused reduced segmentation accuracy for the dHCP pipeline. The dHCP is also computationally demanding, requiring several hours to segment a single infant MRI scan even with high-performance computing. In contrast, ID-Seg takes only 4 seconds per scan. Leveraging deep learning, we offer ID-Seg as a highly efficient, accurate, and reliable approach to segmenting the amygdala and hippocampus in the infant brain.

We focused on the segmentation of the hippocampus and amygdala and found moderate negative associations between the volumes and shapes of these structures and parent-reported behavioral problems at age 2. These negative associations suggest that smaller hippocampus and amygdala at birth relate to greater behavioral problems at age 2. These findings, albeit in a medium sample (n=33), indicate that quantifying morphometrics of limbic substrates may offer important insights into subsequent psychiatric impairment. If replicated in a larger sample, this approach could, for example, but used to help identify infants at high risk for psychiatric impairment. With that knowledge, early preventive strategies and related resources could be focused on those most in need. It is interesting to note that only one prior study reported a significant relationship between morphometrics of the hippocampus and amygdala and CBCL externalizing scores in 12-year-old teenagers (Hanson et al., 2015). Consistent with our findings, Hanson et al. found that smaller hippocampus and amygdala volumes were associated with greater behavioral problems for 12-year-old children.

In this work, the accuracy and reliability of ID-Seg are very promising. However, it is important to consider potential limitations. First, ID-Seg has not been tested in infants of different ages (e.g., 4-6 months). Second, ID-Seg has not been tested on T1w MRI scans. Third, the ground-truth manually segmented dataset, against which we determined the accuracy of ID-Seg was still small. Future work will need to continue developing ID-Seg (e.g., add adaptive domain blocks) and expand the ground-truth manual datasets to include infants of different ages and other subcortical regions (e.g., caudate, putamen). Fourth, in the proof-of-concept analysis, we found that morphometric measures derived from ID-Seg provided strong brain-behavior associations; however, this was based on a relatively small sample. These brain-behavior findings need to be replicated in larger, independent samples.

In sum, we developed a reliable and efficient infant deep learning segmentation framework (**ID-Seg**) to significantly improve the segmentation accuracy of hippocampi and amygdalae in newborn infants. We also demonstrated that **ID-Seg**-derived morphometric measures would provide stronger brain-behavior associations. ID-Seg – an accurate, reliable, and time-efficient segmentation method for infant MRI scans, will improve our capacity to efficiently measure MRI-based brain features relevant to neuropsychological development and ultimately advance the success of quantitative analyses with large-scale datasets.

## Supporting information

Supplemental Materials about Manual Segmentation Protocol

## Acknowledgments

Research reported in this publication was supported by the Environmental influences on Child Health Outcomes (ECHO) program, Office of The Director, National Institutes of Health, under Award Numbers U2COD023375 (Coordinating Center), U24OD023382 (Data Analysis Center), U24OD023319 (PRO Core), and UH3OD023328 (Duarte C, Monk CE, Canino G, and Posner J). The authors wish to thank our ECHO colleagues, the medical, nursing and program staff, as well as the children and families participating in the ECHO cohorts.

