## Supplemental Table and Supplemental Method for "ID-Seg: An Accurate and Reliable Infant Deep learning Segmentation Framework for Limbic Structures"

### Manual Segmentation Protocols Using ITK-SNAP: Hippocampus and Amygdala

#### Part 1: Setting up your tools

##### Loading your files

1. Open your main image in ITK-SNAP.
2. Choose “Add Another Segmentation” and create a new segmentation file.

##### Preparing to label

4. On the main toolbar, select the fourth icon on the first row (paintbrush inspector). For brush style, select the square. For brush size, choose “1.” For brush options, check “3D” and “Isotropic.”
5. On the main toolbar, select the fifth icon that looks like a paint palette on the second row (segmentation label editor). Under available labels, delete unneeded labels and rename labels 1 through 4 as follows:

| Label number | Label name | Label color |
|---|---|---|
| 1 | Right Hippo | red (R:255, G:0, B:0) |
| 2 | Left Hippo | green (R:0, G:255, B:0) |
| 3 | Right Amyg | blue (R:0, G:0, B:255) |
| 4 | Left Amyg | yellow (R:255, G:255, B:0) |

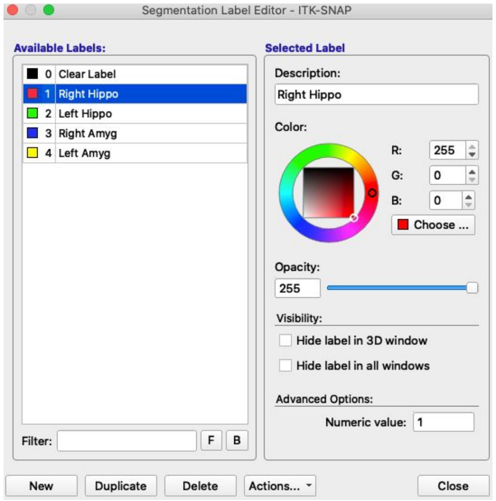


6. On the main toolbar, select the third icon on the second row (image layer inspector). The left hand bar features every layer you’ve loaded. This is where you will toggle between the main image and segmentation layers. Make sure you have selected your segmentation layer before beginning to label.
7. Remain in the image layer inspector. Choose the second tab (contrast). In the Linear Contract Adjustment box, select “Auto.” This should brighten up your image. If you want to toggle with the Curve-Based Contrast Adjustment to change this, feel free to do so.

#### Part 2: Finding where to begin

1. Begin center mass on the axial view (top left of the four images). By "center mass," I mean that your "slice" should land somewhere near the middle of all slides where the hippocampus and amygdala are visible. For example, Figure 1 shows slice 117 of 256 on the axial view of this infant's MRI scan. Figure 2 shows the same slice after labeling. You may need to scroll a bit to find the most visible slice on which to begin. **The idea here is to begin in a place with maximum visibility.** The clearer your first slice, the easier it will be to work from that slice.

**Figure 1:** Slice 117 of 256. unlabeled.

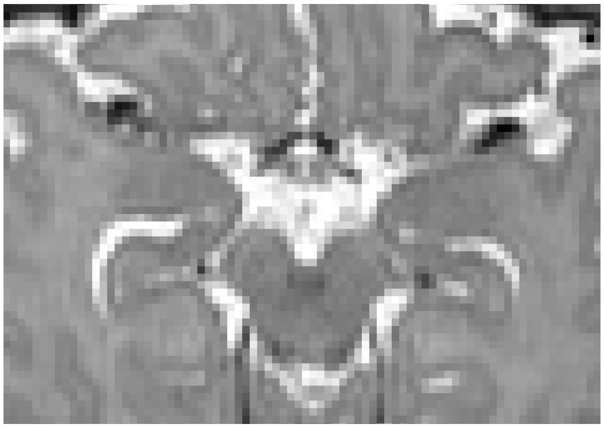

**Figure 2:** Slice 117 of 256. Labelled.

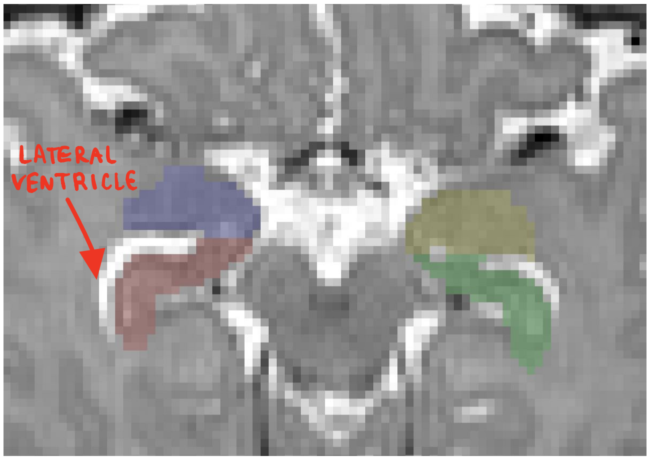

2. You will not need to locate where to begin again until you finish all labels on the axial view. Once you do so, you will need to locate a beginning slice for each of the hippocampus and amygdala pairs on the sagittal view. Since you've already labeled the axial view, you will be able to see those labels translated over to the sagittal view (Figures 3 and 4). Choose a slice where the hippocampus and amygdala are separated by the **temporal horn of the lateral ventricle** (disconnected) AND the hippocampus extends all the way to the **lateral ventricle**. This slice is often about 40 slices away from either the beginning or end (Left: 41 away from 0; Right: 42 away from 130).

**Figure 3**: Slice 41 of 130. Labelled.

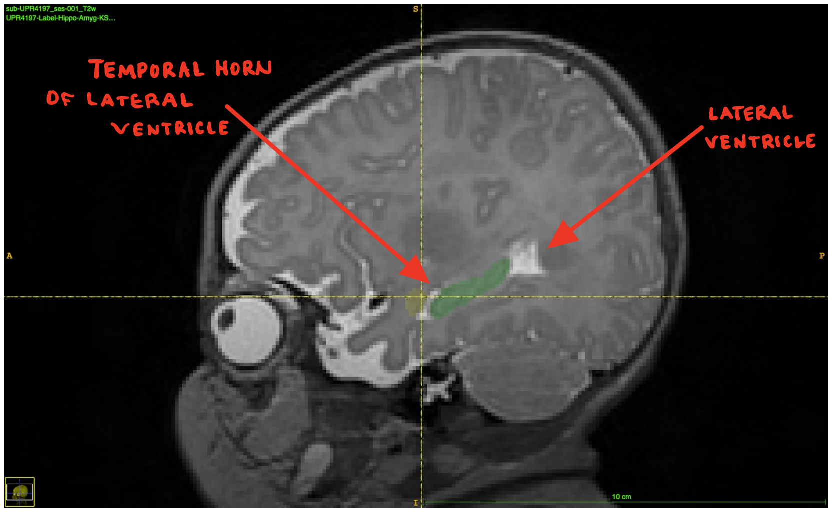


**Figure 4**: Slice 88 of 130. Labelled.

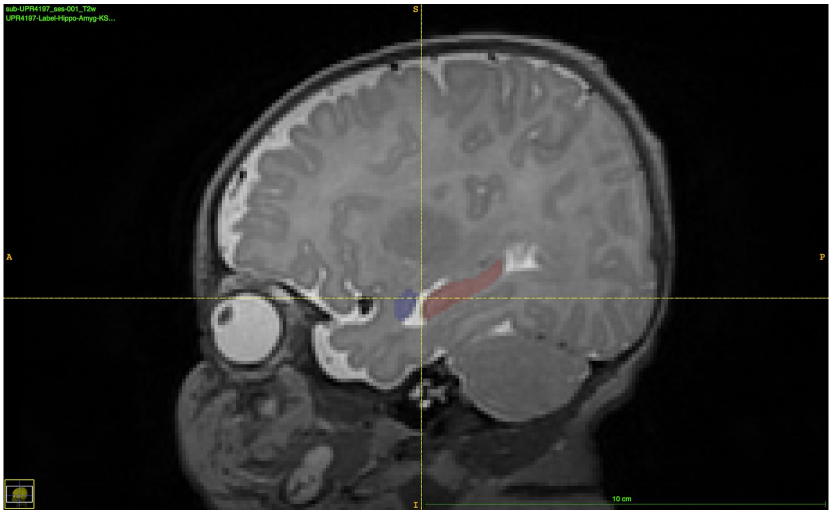

#### Part 3: Establishing an order

Now that you've found your starting point, it's important to know the order in which you'll label.

Segment all labels on the axial view first. Use the following order:

1. Right hippocampus (Label 1)
2. Right amygdala (Label 3)
3. Left hippocampus (Label 2)
4. Left amygdala (Label 4)

Next, repeat the same order in examining the sagittal view. Finally, repeat the process one more time on the axial view to do final touch-ups and edits.

*Note.* Distinguish between right and left by using the markers on ITK-SNAP, not by using your own directionality. The directions are reversed.

### Part 4: Performing Segmentation: Establishing boundaries using landmarks

There are 8 key moments of segmentation on the axial view and 6 key moments on the sagittal view. I will walk you through all 14. By using these pivotal shifts in segmentation protocol as your baseline, you'll be able to fill in the surrounding slices easily. **Pay attention to the hand-drawn annotations. Those will guide you in understanding the landmarks you will use as boundaries for labelling.** For the purposes of our segmentation, we will disregard the coronal view.

#### Axial view segmentation

1. Start where you saw the most visibility in step 1 of part 2: finding where to begin. This will be your simplest segmentation image.

Slice 117 of 256: Previously-established starting point (unlabeled, labeled)

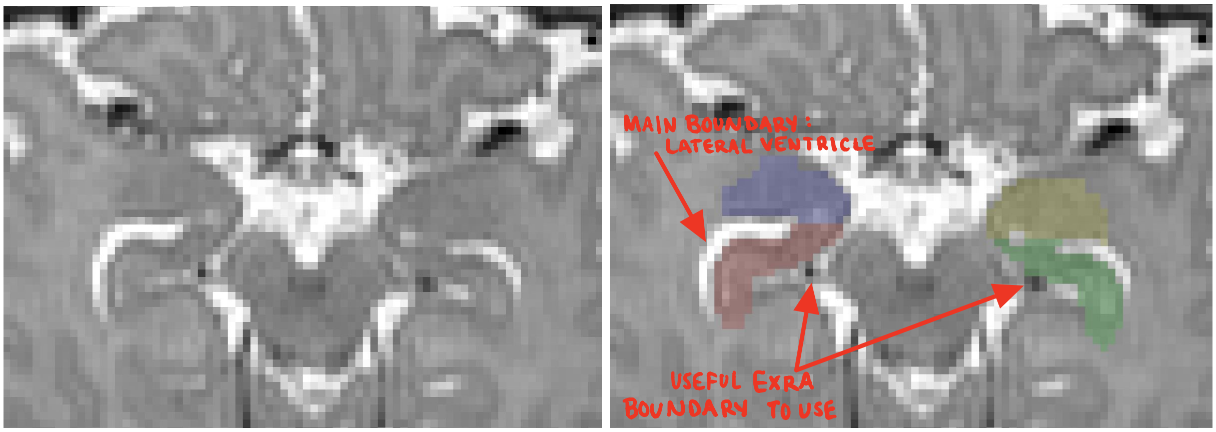


2. Scroll down until you begin to see the hippocampus and amygdala "splitting," or no longer touching. When this happens, the amygdala should remain around where it was on the image, right next to the white matter that lines each thalamus. Meanwhile, the hippocampus should follow the boundary established by the white matter to its left (see images below).
    a. The amygdala should remain around where it was on the image, right next to the white matter that lines each thalamus.
    b. The hippocampus should follow the boundary established by the white matter to its left (see image).
    c. Make sure that all hippocampal labeling is connected to itself. This means that, once the hippocampus and amygdala disconnect, no hippocampus labeling should remain attached to the amygdala. See image for clarification.

*Note.* This "splitting" may occur at different rates on each side—it's important to note that the right and left sides are NOT mirrored. They are often different in shape and thickness. For example, the image below shows that the left hippocampus and amygdala still connect (since there is gray matter of equal shading connecting them), but the right hippocampus and amygdala don't connect.

Slice 120 of 256: One hippocampus "splits" from its amygdala, while the other remains thinly connected (unlabeled, labeled)

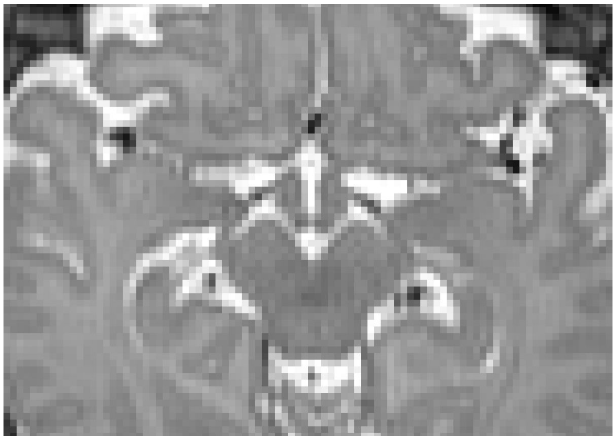

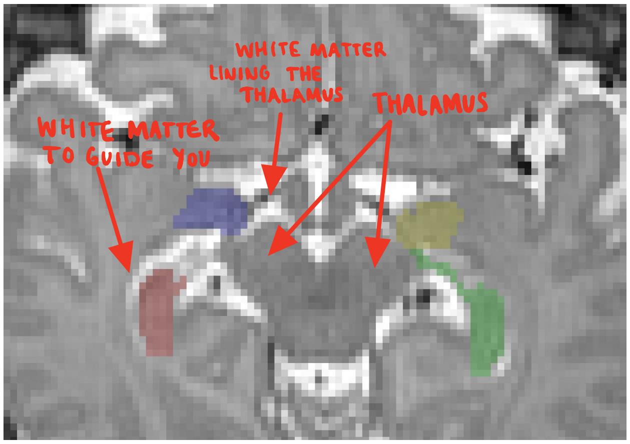


3. Continue scrolling down. The amygdala will disappear from the images. You can tell by tracking the dark gray matter. Once it is no longer visible, there's no more amygdala to label. Use the labelled boundaries to contain your hippocampal labeling:

Slice 123 of 256: Use differences in shade to distinguish between the hippocampi and their boundaries. There is white matter that lines the bottom of the hippocampus and almost black-looking matter that lines the top (unlabeled, labeled)

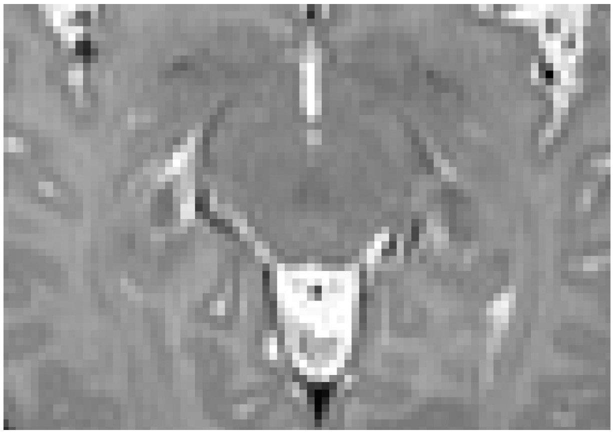

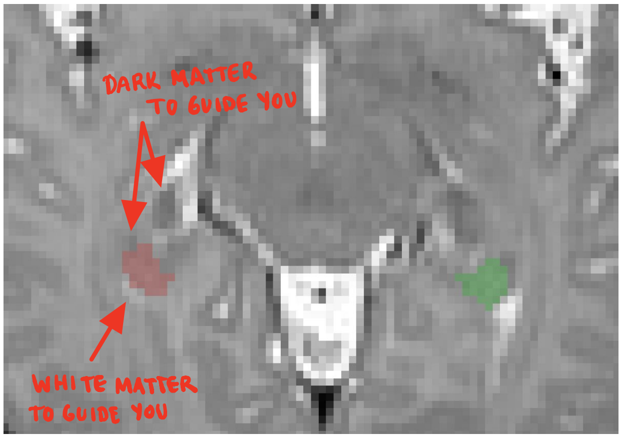

4. After that, scroll back up. You'll pass your starting point and continue until the hippocampus no longer curves down like seahorse. The hippocampus and amygdala are connected on a wider portion now.
    a. You'll use the **lateral ventricle** (labelled) as your boundary for splitting the amygdala and hippocampus.
    b. Your amygdala should look similar to a mushroom top in shape.
    c. Finally, check that your splitting point results in about ⅔ hippocampus and ⅓ amygdala. The hippocampus shouldn't be dramatically bigger than the amygdala at this point. Since the hippocampus and amygdala have similarly colored gray matter, we use this rule to keep segmentations standardized.

Slice 114 of 256: The lateral ventricle serves as a boundary between the hippocampus and amygdala (unlabeled, labeled)

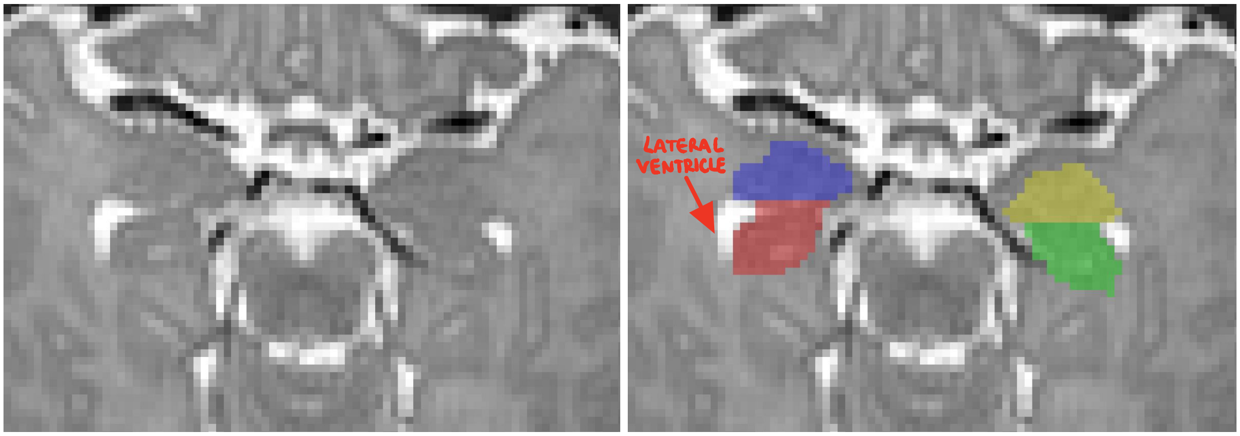


5. Finally, find where the hippocampus and amygdala split again. As the structures get smaller, continue to use the lateral ventricle as your boundary, but split the hippocampus and amygdala down the middle (so each are similar sizes now). Here's a series of images describing before and after the final split:

Slice 111 of 256: Before the hippocampus and amygdala split (unlabeled, labeled)

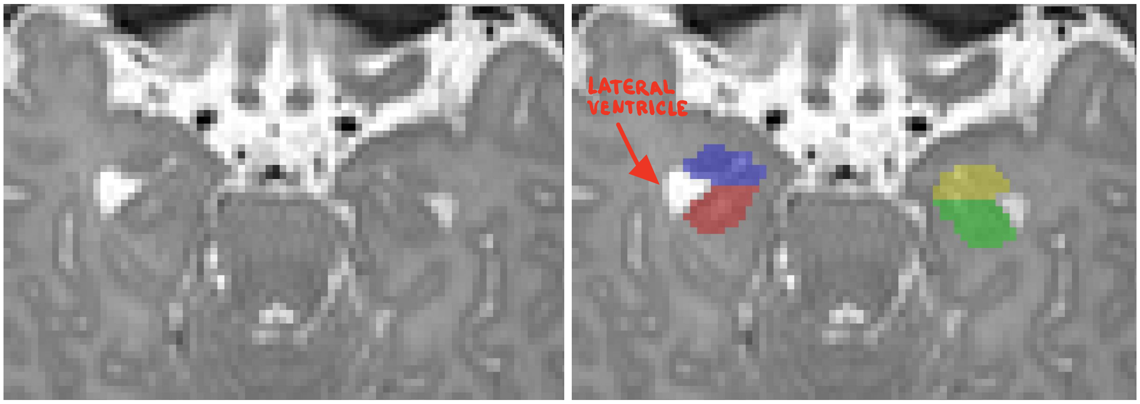

Slice 110 of 256: After the hippocampus and amygdala split, use the lateral ventricle to separate them (unlabeled, labeled)

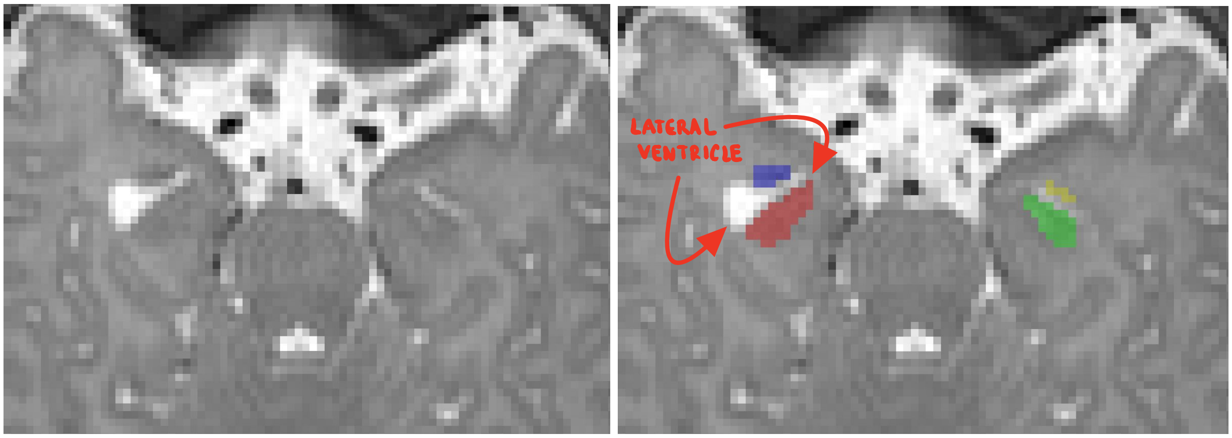


6. This is nearly the end of your image. The amygdala will disappear as you continue scrolling up, and the hippocampus will shrink. Just follow the same color of dark gray matter until that color is no longer visible. Another good way to know when to stop is when you can no longer see the lateral ventricle.

#### Sagittal view segmentation

1. Begin at the image you determined as your starting point. Use the lateral ventricle as your guide.

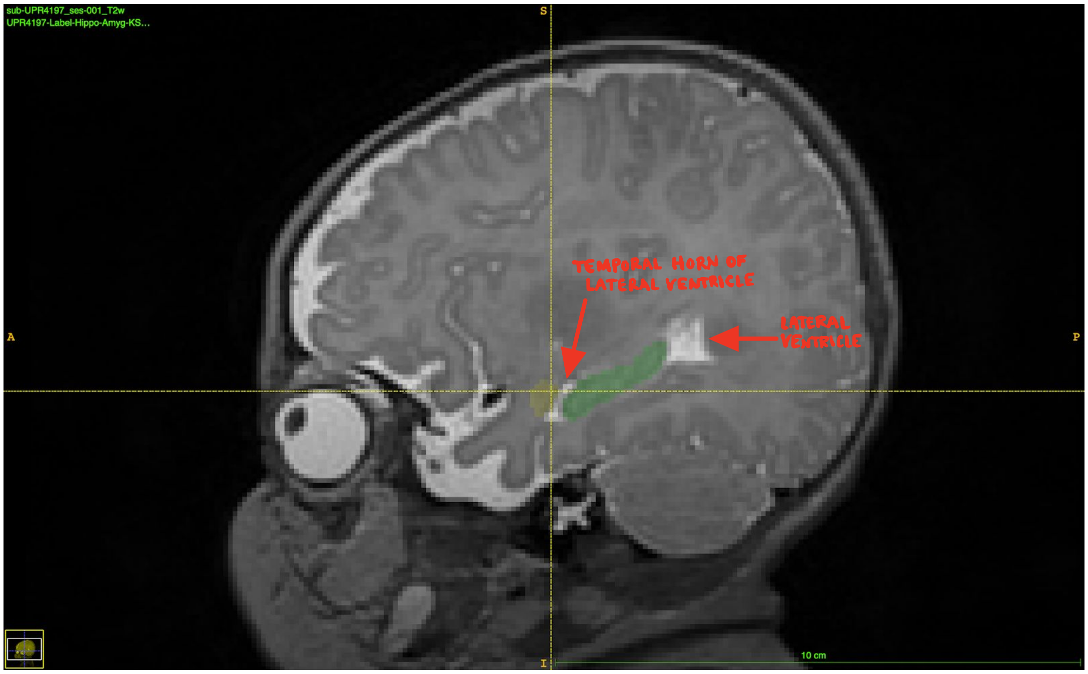

2. Find the end boundary.

Slice 38 of 130: Final slice before the end boundary of the hippocampus (unlabeled, labeled)

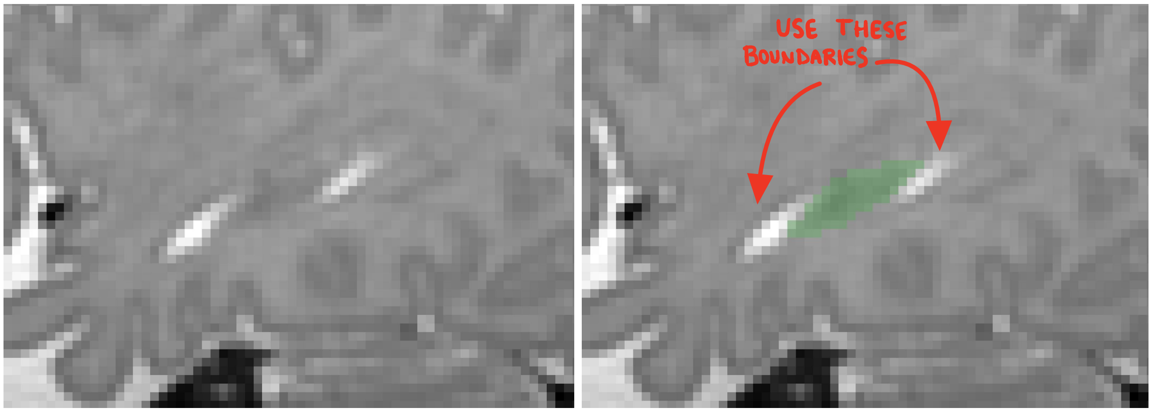


Slice 37 of 130: No hippocampus to label. Notice the lack of dark gray matter that was present in slice 38, which signalled the presence of the hippocampus.

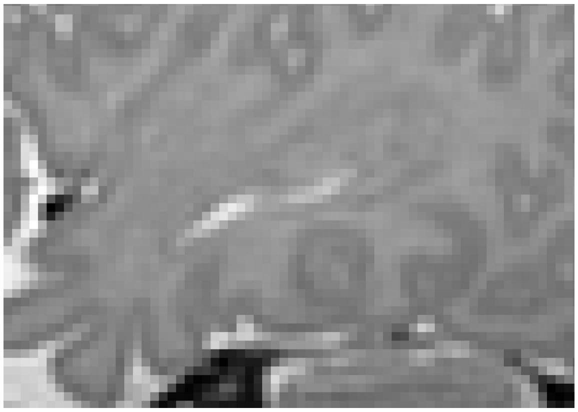

3. Find where the hippocampus and amygdala connect.

Slice 41 of 130: Hippocampus and amygdala do not connect (unlabeled, labeled)

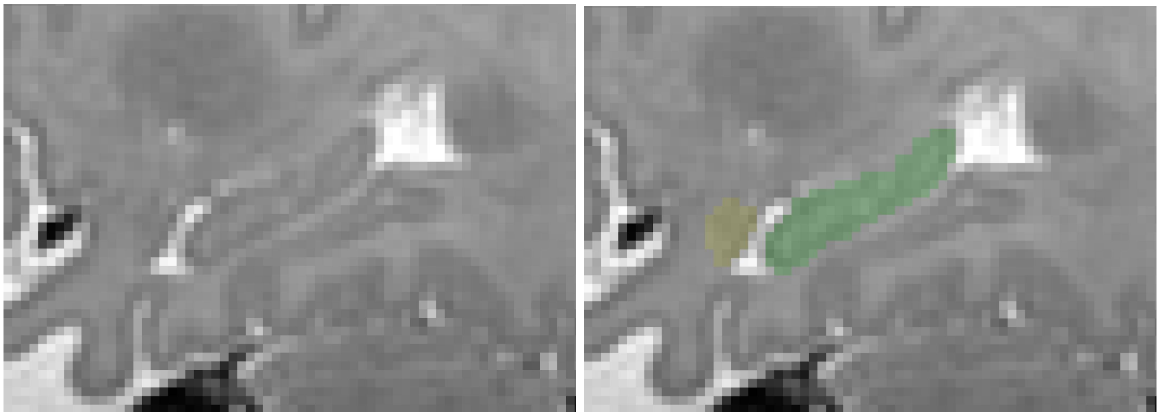

Slice 40 of 130: Hippocampus and amygdala connect (unlabeled, labeled)

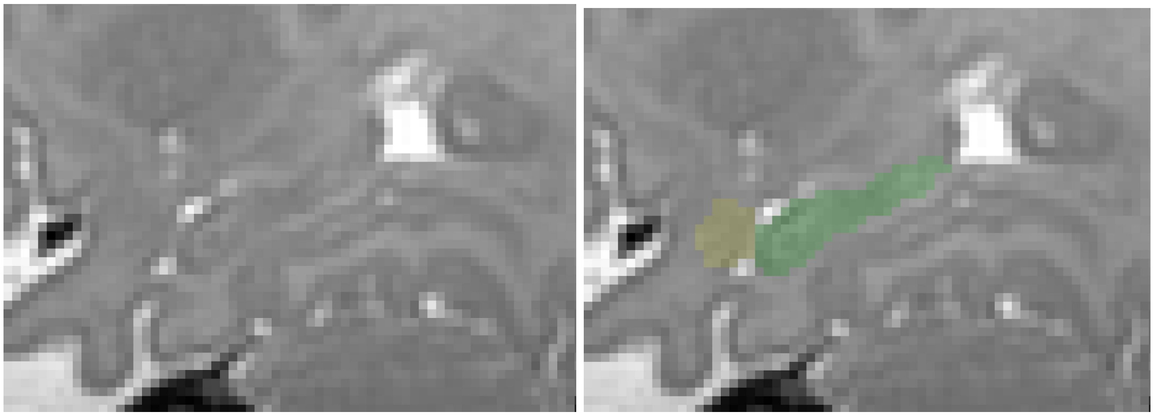

4. Find where the tail ends

Slice 42 of 130: Tail extends to the fornix and lateral ventricle (unlabeled, labeled)

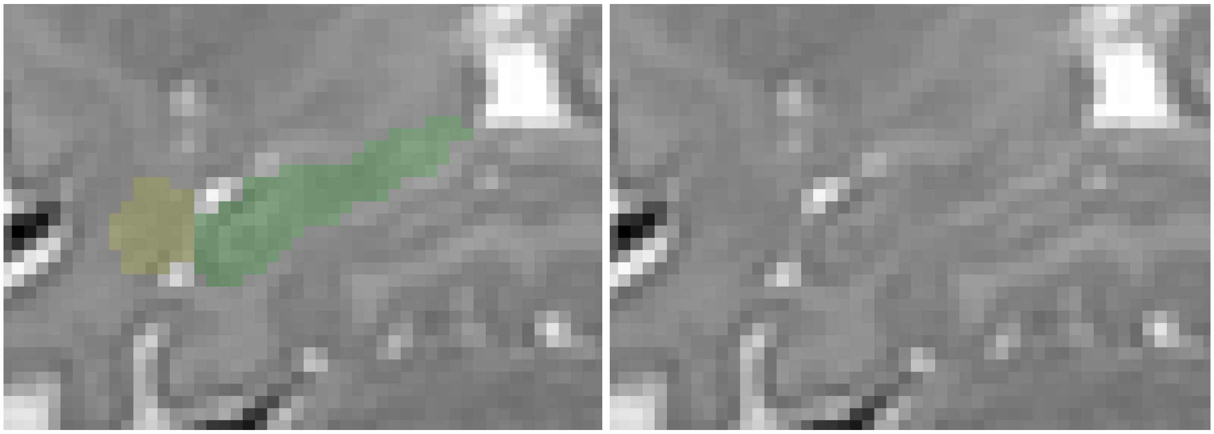

Slice 43 of 130: Dark gray matter signals that there is no more tail (unlabeled, labeled)

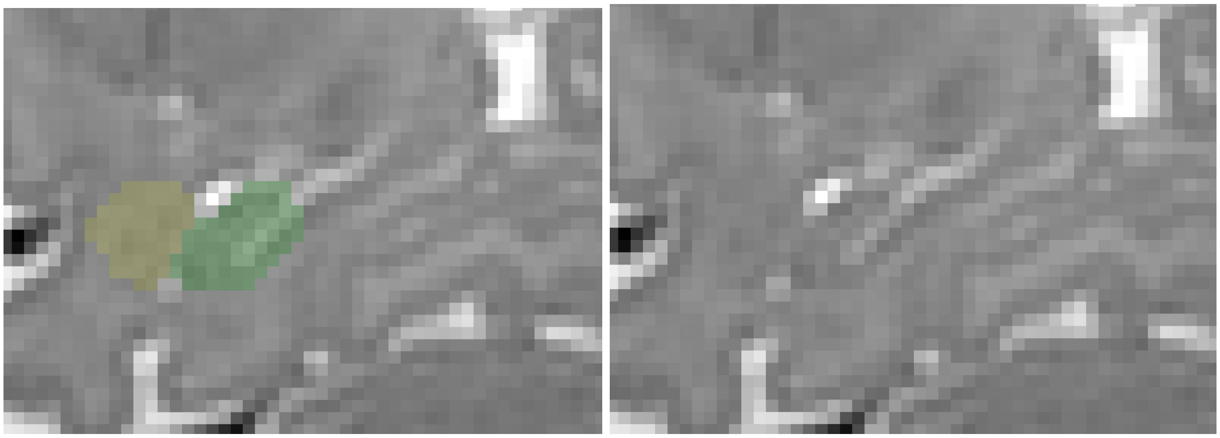

5. Use boundaries to your advantage.

Slice 51 of 130: Splitting the hippocampus and amygdala using white matter as a halfway point (unlabeled, labeled)

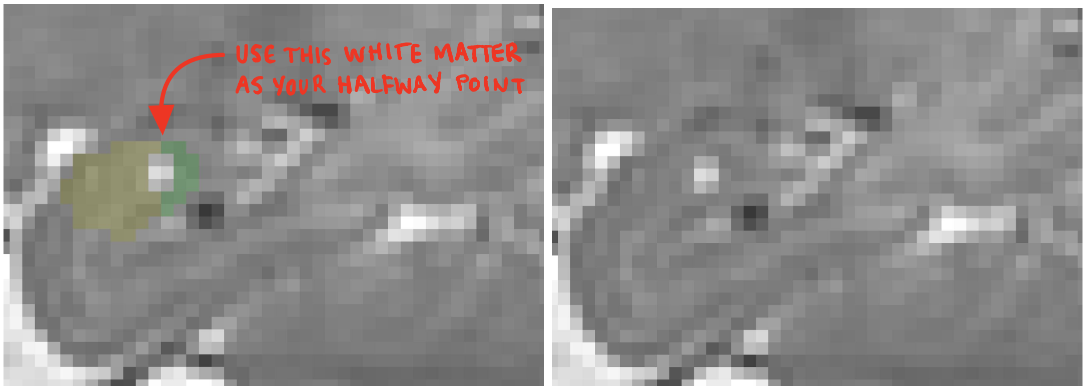

6. End with the amygdala.

Slice 54 of 130: The final traces of the hippocampus (unlabeled, labeled)

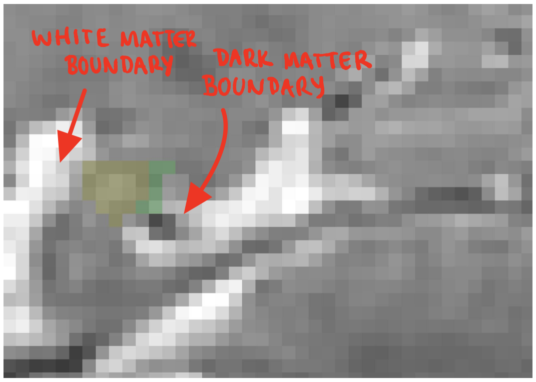


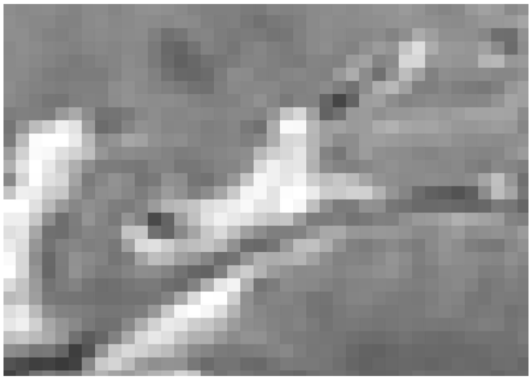

Slice 55 of 130: Amygdala only (unlabeled, labeled)

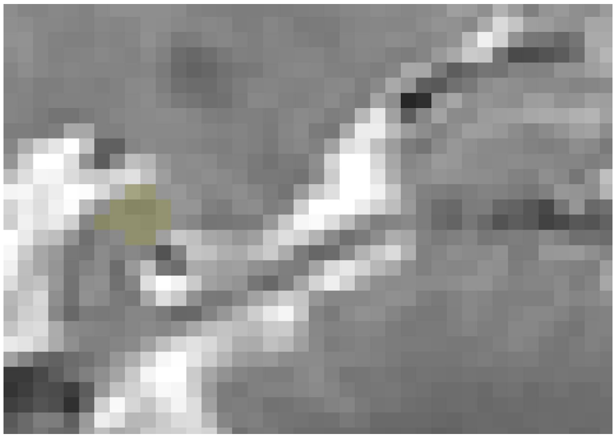

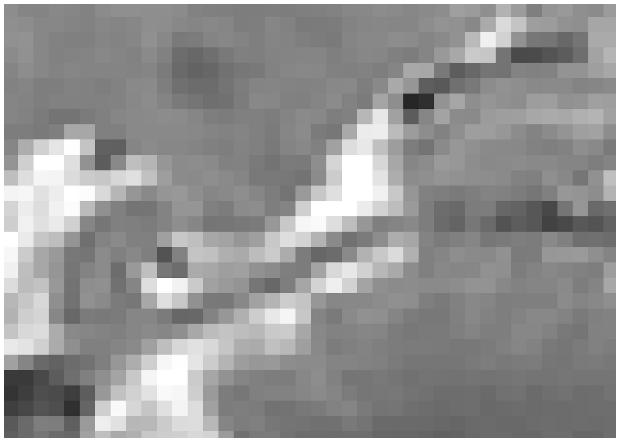

#### Part 5: Reviewing your work

1. Update your 3D image. Do the structures look reasonable in size and shape? Are there any protruding spikes? Spikes indicate areas that are likely mistakes. Go back and check suspect areas.
2. Make sure that there are no “floating voxels.” Your labels should all be connected to each other. It takes time, but it’s crucial to double check that your work is neat.
3. Toggle the label view on and off. Reevaluate your labeling. Do your eyes still see the shapes the same way? If not, edit boundaries accordingly.
4. Ensure that you haven’t labeled the lateral ventricle or any other surrounding white matter.
5. Do a quick scan through the axial view by scrolling through all slices. Do the slices look cohesive, or are they inconsistent? Edit accordingly.

#### Part 6: Saving your work

1. Go to the main toolbar and click the third icon on the second row (Image Layer Inspector).
2. On the left hand bar, click the player with YOUR labels.
3. Click the small arrow in the corner of that layer and choose “save image.” Use the following format to name your segmentation file: [File Name]-Label-Hippo-Amyg-[Your Initials]-[Date].nii.gz
