## Supplemental Materials about Manual Segmentation Protocol for "ID-Seg: An Accurate and Reliable Infant Deep learning Segmentation Framework for Limbic Structures"

### Supplemental Materials: Datasets

This study was a secondary data analysis from infants with MRI scans acquired as part of four neurodevelopmental studies.

**Study 1 *Environmental influences on child health outcomes(ECHO) – Boricua Youth Study (BYS) at the New York State Psychiatric Institute (ECHO-BYS-NYSPI)* –** Enrollment began in 2016 and is ongoing. The sample comes from the New York City site for the Boricua Youth Study (BYS-NY), an epidemiological study of Puerto Rican families from the South Bronx, New York. Pregnant women within the BYS-NY cohort (or pregnant women whose partner was part of BYS-NY) were recruited to participate. The sample included healthy pregnant women, ages 18–45. The New York State Psychiatric Institute Institutional Review Board (NYSPI-IRB) approved all the procedures and participants provided informed consent. Images were acquired on a GE MR750 3T whole-body scanner with an in-vivo 8-channel head coil and a GE 3T Signa Premier whole-body scanner with a GE 48-channel head coil.

***Study 2 ECHO-BYS at the University of Puerto Rico (ECHO-BYS-UPR )* –** Enrollment began in 2016 and is ongoing. The sample comes from the San Juan, Puerto Rico site for the Boricua Youth Study (BYS-PR). Enrollment followed the same criteria as in Study 1**.** UPR-IRB approved all the procedures and participants provided informed consent. Images were acquired on a GE MR750 3T whole-body scanner with an in-vivo 8-channel head coil.

### Supplemental Table 1. Inter-rater Agreement Assessment.

For each structure and infants, the inter-rater agreement of the manual tracing was assessed by Dice Similarity Score (DSC). In the table below, we presented all inter-rater agreements as well as mean and standard deviation (SD). The overall inter-rater reliability is 0.76 (0.06).

| | Rater 1 vs Rater 2 | | | | Rater 1 vs Rater 3 | | | | Rater 2 vs Rater 3 | | | |
|---|---|---|---|---|---|---|---|---|---|---|---|---|
| | R Hippocampus | L Hippocampus | R Amygdala | L Amygdala | R Hippocampus | L Hippocampus | R Amygdala | L Amygdala | R Hippocampus | L Hippocampus | R Amygdala | L Amygdala |
| sub01 | 0.72 | 0.73 | 0.71 | 0.76 | 0.69 | 0.78 | 0.72 | 0.73 | 0.83 | 0.71 | 0.80 | 0.76 |
| sub02 | 0.78 | 0.76 | 0.64 | 0.69 | 0.72 | 0.66 | 0.70 | 0.70 | 0.70 | 0.63 | 0.70 | 0.75 |
| sub03 | 0.73 | 0.75 | 0.76 | 0.79 | 0.73 | 0.66 | 0.78 | 0.78 | 0.77 | 0.66 | 0.77 | 0.78 |
| sub04 | 0.74 | 0.71 | 0.75 | 0.73 | 0.78 | 0.79 | 0.68 | 0.75 | 0.73 | 0.77 | 0.68 | 0.73 |
| sub05 | 0.69 | 0.69 | 0.65 | 0.64 | 0.76 | 0.79 | 0.72 | 0.80 | 0.78 | 0.68 | 0.66 | 0.62 |
| sub06 | 0.65 | 0.75 | 0.66 | 0.63 | 0.66 | 0.74 | 0.67 | 0.74 | 0.74 | 0.73 | 0.70 | 0.68 |
| sub07 | 0.78 | 0.78 | 0.79 | 0.77 | 0.81 | 0.81 | 0.77 | 0.78 | 0.78 | 0.79 | 0.75 | 0.80 |
| sub08 | 0.80 | 0.77 | 0.83 | 0.81 | 0.79 | 0.81 | 0.78 | 0.81 | 0.81 | 0.80 | 0.82 | 0.85 |
| sub09 | 0.76 | 0.75 | 0.60 | 0.73 | 0.81 | 0.79 | 0.67 | 0.72 | 0.82 | 0.74 | 0.75 | 0.73 |
| sub10 | 0.75 | 0.77 | 0.70 | 0.75 | 0.83 | 0.78 | 0.74 | 0.70 | 0.76 | 0.76 | 0.73 | 0.66 |
| sub11 | 0.81 | 0.76 | 0.79 | 0.76 | 0.81 | 0.81 | 0.69 | 0.73 | 0.80 | 0.73 | 0.64 | 0.74 |
| sub12 | 0.73 | 0.74 | 0.69 | 0.77 | 0.81 | 0.76 | 0.77 | 0.79 | 0.71 | 0.68 | 0.69 | 0.76 |
| sub13 | 0.77 | 0.77 | 0.79 | 0.79 | 0.84 | 0.86 | 0.76 | 0.84 | 0.76 | 0.75 | 0.68 | 0.75 |
| sub14 | 0.85 | 0.82 | 0.80 | 0.78 | 0.88 | 0.82 | 0.81 | 0.80 | 0.85 | 0.81 | 0.78 | 0.72 |
| sub15 | 0.78 | 0.80 | 0.72 | 0.70 | 0.83 | 0.77 | 0.79 | 0.79 | 0.77 | 0.75 | 0.72 | 0.69 |
| sub16 | 0.81 | 0.84 | 0.73 | 0.61 | 0.85 | 0.85 | 0.83 | 0.82 | 0.79 | 0.81 | 0.74 | 0.66 |
| sub17 | 0.76 | 0.77 | 0.72 | 0.82 | 0.84 | 0.85 | 0.76 | 0.83 | 0.75 | 0.78 | 0.71 | 0.76 |
| sub18 | 0.81 | 0.78 | 0.71 | 0.71 | 0.82 | 0.83 | 0.78 | 0.81 | 0.80 | 0.78 | 0.76 | 0.66 |
| sub19 | 0.78 | 0.77 | 0.74 | 0.78 | 0.87 | 0.84 | 0.80 | 0.81 | 0.81 | 0.80 | 0.73 | 0.72 |
| sub20 | 0.81 | 0.75 | 0.77 | 0.71 | 0.84 | 0.84 | 0.71 | 0.72 | 0.76 | 0.79 | 0.69 | 0.65 |
| **Mean** | 0.76 | 0.76 | 0.73 | 0.74 | 0.80 | 0.79 | 0.75 | 0.77 | 0.78 | 0.75 | 0.72 | 0.72 |
| **SD** | 0.05 | 0.03 | 0.06 | 0.06 | 0.06 | 0.06 | 0.05 | 0.05 | 0.04 | 0.05 | 0.05 | 0.06 |

**Supplemental Figure 1. Multilinearity Check**
Before the multivariate analysis, we examined the collinearity among variables by calculating variance inflation factor (VIF). Conventionally, VIF less than 5 indicates a low correlation of that predictor with other predictors. A value between 5 and 10 indicates a moderate correlation, while VIF values larger than 10 are a sign for high (*James et al. 2013*). In figure below, 5 out of 8 brain metrics from R/L hippocampus and amygdala has shown moderate-to-high correlations.

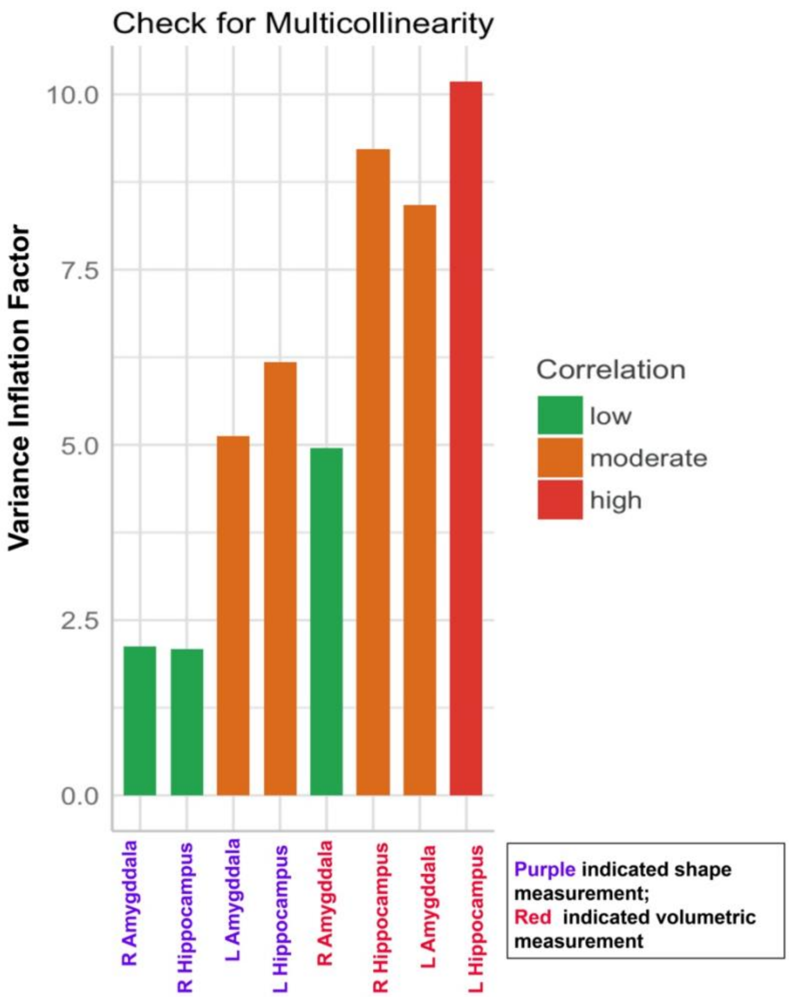
